# ME-ACP: Multi-view Neural Networks with Ensemble Model for Identification of Anticancer Peptides

**DOI:** 10.1101/2021.11.22.469543

**Authors:** Guanwen Feng, Hang Yao, Chaoneng Li, Ruyi Liu, Rungen Huang, Xiaopeng Fan, Ruiquan Ge, Qiguang Miao

## Abstract

Cancer remains one of the most threatening diseases, which kills millions of lives every year. As a promising perspective for cancer treatments, anticancer peptides (ACPs) overcome a lot of disadvantages of traditional treatments. However, it is time-consuming and expensive to identify ACPs through conventional experiments. Hence, it is urgent and necessary to develop highly effective approaches to accurately identify ACPs in large amounts of protein sequences. In this work, we proposed a novel and effective method named ME-ACP which employed multi-view neural networks with ensemble model to identify ACPs. Firstly, we employed residue level and peptide level features preliminarily with ensemble models based on lightGBMs. Then, the outputs of lightGBM classifiers were fed into a hybrid deep neural network (HDNN) to identify ACPs. The experiments on independent test datasets demonstrated that ME-ACP achieved competitive performance on common evaluation metrics.

## 1 Introduction

Cancer has evolved into one of the serious diseases which threaten human life. According to the report from the world health organization, various forms of cancer cause millions of deaths every year, and the number is increasing year by year. Therefore, how to find effective methods to treat cancer is an urgent problem. Traditional treatments often do not work well such as radiation therapy, targeted therapy and chemotherapy [1]. So, more effective treatments are urgent needed to overcome the lots of aforesaid disadvantages.

In recent years, the peptide-based therapy offers a promising method for cancer treatments. This method can usually kill cancer cells in a targeted way without affecting the function of the normal cells. ACPs with a less length than 50 amino acids show the prominent advantages compared with the traditional cancer treatments. And they display high selectivity to kill the cancer cells. While most ACPs are cationic, cancer cell membranes are generally anionic, which allow the anticancer peptides to target and kill cancer cells through electrostatic interactions [2, 3]. However, it is time-consuming and arduous to do experiments to identify the potential ACPs from the huge protein databases. Artificial intelligence methods are suitable for the detection task of ACPs as the alternative methods. They can be used to screen the protein sequences whether they have the anticancer function through machine learning, deep learning and other methods, so as to shorten the experimental period and improve the efficiency of cancer treatment.

Nowadays, the accurate prediction of ACPs has become a hot issue in the field of computational biology based on machine learning methods. In general, most of the methods mainly contain two key technologies. One is effective feature representation of protein sequences, the other is powerful machine learning algorithm. For feature representation, if the feature engineering of the sequence is done well, it will make a good foreshadowing for the following classification. The first predicted tool named antiCP encodes proteins as binary features and feeds them into the support vector machine (SVM) classifier to identify the ACPs[4]. Chou’s pseudo acid amino composition (PseAAC) can effectively improve the prediction accuracy of ACPs[5]. A variety of feature extraction methods from the information of amino acids were integrated in ACPP[6]. Li et al. improved the accuracy of prediction by the feature’s fusion[7]. MLACP employed SVM and random forest (RF) classifiers to identify ACPs using the features extracted from the amino acid sequence[8]. ACPred-FL fed several types of features extracted from the sequences into an SVM classifier to get the first predicted result. Then a two-step feature selection method was employed to find the informative feature representation[9]. ACPred-Fuse embedded the class and probabilistic multi-view information into the ACPs prediction[10]. More recently, deep learning techniques have also been explored to predict ACPs. ACP-DL fed the features from the k-mer sparse matrix and one-hot encoding into the long short-term memory (LSTM) model to predict ACPs[11]. DRACP fed the extracted 56 features from composition into deep belief network to reduce the dimension of features. Then random relevance vector machines were used to identify ACPs[12]. DeepACPpred employed a hybrid CNN-RNN architecture to predict ACPs[13]. iACP-DRLF obtained the features through two pretrained deep representation learning embedding models. Then, the ensemble classifiers were used to find the optimized solution[14]. ACPred-LAF can auto extract features based on multisense and multiscaled embedding[15]. GRCI-Net employed a reduced fusion feature through principal component analysis. Then it integrated a bidirectional long- and short-term memory (Bi-LSTM) and a multiscale dilated convolution network to predict the anticancer peptides[16] In terms of initial feature extraction, protein sequence encoding was used by most current methods, which extracted features from entire protein sequence. But many of them will lack local information of each residue. Amino acid coding has been neglected in the past. Furthermore, some correlations between residuals were difficult to find for the extracted features.

In this work, we proposed a Multi-view neural networks with Ensemble model (ME-ACP) to identify anticancer peptides accurately and effectively. The model architecture was shown in Fig.1. Residue level and peptide level feature representation methods were employed to extract the initial features. And then, the generated features were fed into multiple lightGBMs[17] respectively to obtain probabilities representation of the samples. The informative probabilities were concatenated as a vector inputting into HDNN, which consisted of residual module[18] and Bi-directional Long Short-Term Memory (Bi-LSTM) module [19]. With the two-path HDNN, ME-ACP could capture the local information and long-range dependency to achieve the integration of multi-view information. Comparative experiments demonstrated the outstanding performance of ME-ACP on the cross-validation and independent datasets.

**Fig.1.**
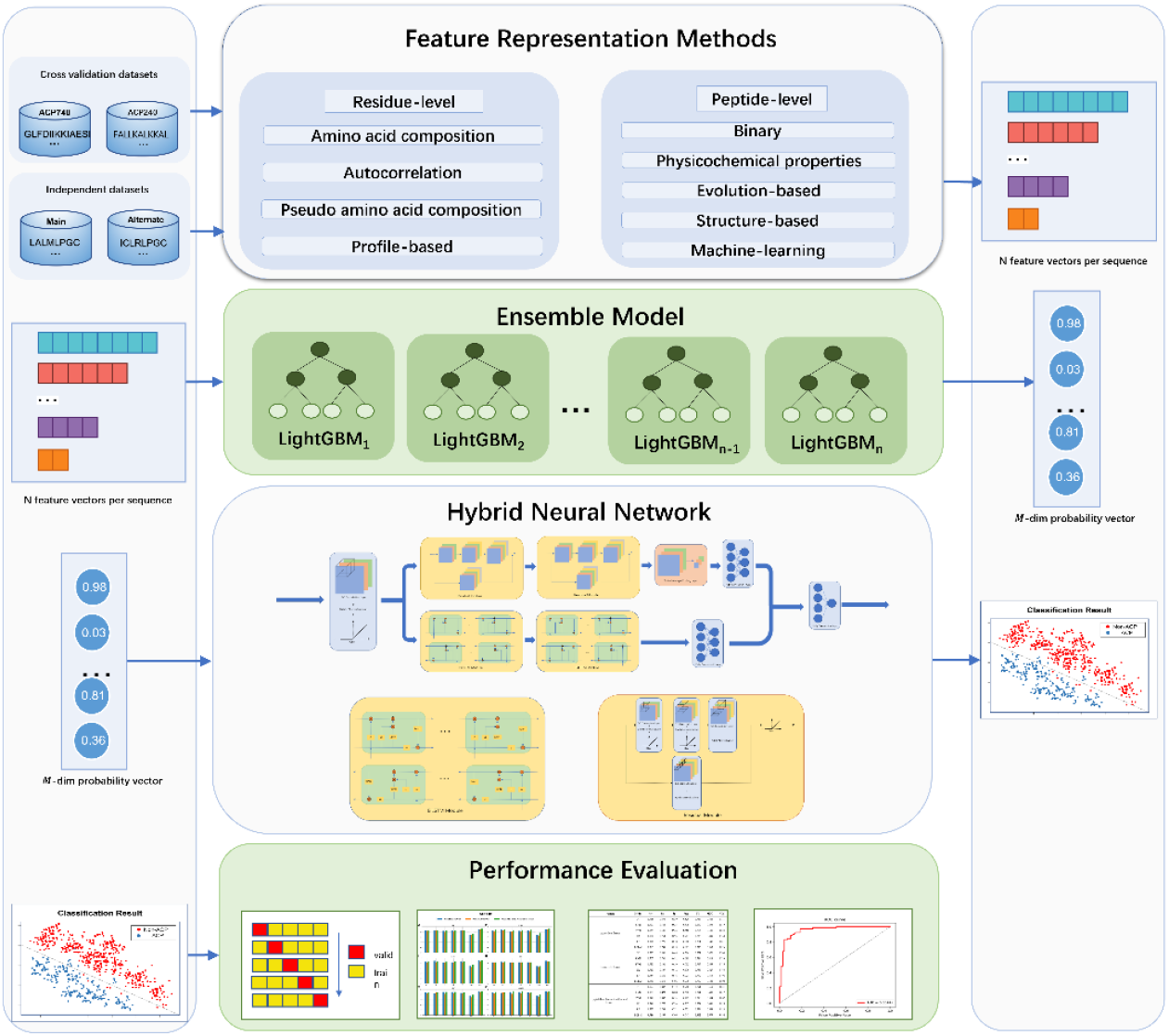
The overview of ME-ACP. Residue level and peptide level features were inputted into multiple lightGBMs. The outputs from the lightGBMs as the low-dimensional features were fed into a hybrid deep neural network, which included a sequential convolution module followed by two parallel paths of residual module and Bi-LSTM module, with global average pooling (GAP) followed by full connected layer (FC) and FC respectively. Finally, FC with sigmoid activation function was employed to identify ACPs and no-ACPs.

## 2 Materials and methods

### 2.1 Datasets

Two groups of updated datasets were applied to evaluate the ME-ACP performance. For them, one was from ACP-DL and the other was from Anticp-2.0[20] which were used for cross-validation and independent experiments, respectively. The cross-validation datasets were constructed in ACP-DL which included ACP740 and ACP240. ACP740 contained 376 ACPs and 374 non-ACPs. The ACP240 consisted of 129 experimentally validated ACPs and 111 AMPs as non-ACPs. There were low redundancy and no overlap in these two datasets. The independent datasets from Anticp-2.0 were composed of Main and Alternate datasets[14]. The main diversity between the two datasets was different sources of negative samples. Main dataset contained 861 experimentally validated ACPs and 861 non-ACPs. The negative samples in the Main dataset were all AMPs without anticancer activity. Alternate dataset consisted of 970 experimentally validated ACPs and 970 non-ACPs. The negative samples were randomly extracted from SwissProt[21].

**Table 1.** Dataset distribution

| Dataset | Cross Validation |  |  |  |
| --- | --- | --- | --- | --- |
|  | ACP |  | Non-ACPs |  |
| ACP-740 | 376 |  | 374 |  |
| ACP-240 | 129 |  | 111 |  |
| Dataset | Independent test |  |  |  |
|  | train-ACP | train-non-ACP | test-ACP | test-non-ACP |
| Main | 689 | 689 | 172 | 172 |
| Alit | 776 | 776 | 194 | 194 |

### 2.2 Feature representation

In general, protein sequences are represented with digital vector as a natural language by using twenty English letters. The digital representation of protein sequences is divided into two categories: residue encoding scheme and peptide encoding scheme. The peptide encoding descripts the global information of sequence but the residue-specific information is lost. The residue encoding scheme ignores the sequential information. Therefore, the combination of residue level and peptide level encoding scheme can fully model sequence information locally and globally.

The features extraction methods on residue level were classified into five categories: binary, physicochemical properties, evolution-based, structure-based and machine-learning. The protein sequences were represented by 19 feature extraction methods in this work. The features extraction methods on peptide level were divided into four categories: amino acid composition, autocorrelation, pseudo amino acid composition and profile-based features.

First, residue level and peptide level features were extracted by a diversity of feature descriptors. For the feature representation scheme on peptide level, entire peptide sequence *s* was converted into a fixed dimension feature vector *F*_*p*_ through each function *g* in *G*, which is a set including different feature extraction methods on peptide level.

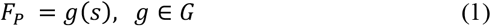

For the feature representation scheme on residue level, each residue was represented as a vector *f*_*i*_ in each protein sequence. Multiple feature vectors on single residue were concatenated into a sequence feature vector *F*_*R*_.

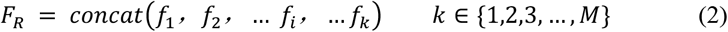

*F*_*R*_ is the sequence feature vector of N-terminus or C-terminus, where *f*_*i*_ is a feature vector of *ith* residue in the sequence. *M* is the minimum length of the peptide sequences in datasets.

In order to improve the expression of features, a few of classic machine learning model was employed to handle the sequence feature extracted by a variety of methods. As a matter of convenience, peptide level feature vector *F*_*P*_ and residue level feature vector *F*_*R*_ were represented by *x*_*i*_. The *x*_*i*_ was fed into machine learning model 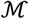, such as the light gradient boosting machine (lightGBM), random forest (RF), decision trees (DT), k-nearest neighbor (KNN), support vector machines (SVM), logistic regression (LR), to obtain a probability *p*_*i*_.

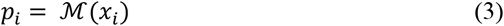

Where *p*_*i*_ ∈ ℝ^1^, *i* = 0, 1 … *M*.

Through the processing of traditional machine learning, high-dimensional protein features of different lengths were converted into low-dimensional features of fixed length to alleviate over-fitting problem in HDNN.

### 2.3 Neural network architecture

In neural network architecture part, HDNN was constructed to predict ACPs and non-ACPs from different views. The probability vector *P* ∈ ℝ^*M*^ from lightGBMs was inputted into HDNN.

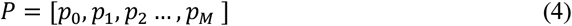

The classic residual structure proposed by the ResNet[18] could extract the local feature information and avoid over-fitting effectively to greatly improve the degradation of the model. So, HDNN utilized the residual module to explore the local information in the protein sequences. Moreover, two layers of Bi-LSTM were considered to capture long range dependency. Therefore, HDNN could capture potential multiple views of sequence information by residual module 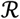 and Bi-LSTM module 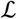.

HDNN started at a sequential convolution module 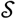 with 1D convolution, batch normalization (BN) and ReLU. With the sequential convolution module 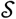, *P* was converted into a 16 × *M* feature.

Two residual convolution layers, which both consisted of three layers of 1D convolution layers and an identity mapping in parallel, obtained the 256 × *M* feature *W* = [*w*_0_, *w*_1_, *w*_2_, …, *w*_256_]^*T*^ from Equation (3).

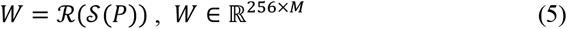

Then, global average pooling (GAP) transformed the 256 × *M* feature into a 256-dimensional vector *V*_1_. In addition, the feature obtained from the sequential convolution module 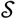 was fed into the Bi-LSTM module, which generated a 128-dimensional vector *V*_2_.

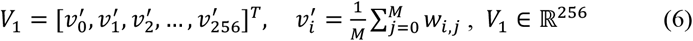

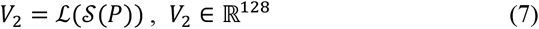

Furthermore, the outputs of residual module and Bi-LSTM module were converted into 64-dimensional vectors by two FCs *F*_1_ and *F*_2_, respectively. Finally, *Pre* from another FC *F* and sigmoid activation function *δ*, were employed to predict ACPs and non-ACPs by serial concatenated two-path 64-dimensional vectors.

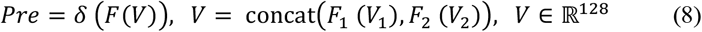

### 2.4 Evaluation

In this study, accuracy (Acc), sensitivity (Sn), specificity (Sp), precision (Prec), F1 score (F1), Matthews correlation coefficient (MCC) and area under receiver operating characteristic curves (AUC) were employed to verify the performance of the model.

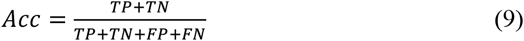

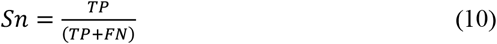

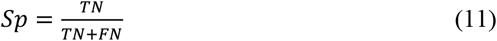

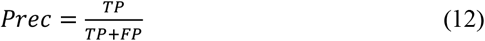

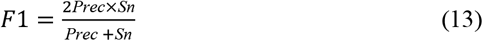

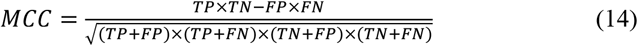

The TP, FP, TN, FN are the brief abbreviations for true positives, false positives, true negatives, and false negatives, respectively.

K-fold cross validation and independent test were used to evaluate the model on different datasets. First, the original dataset was divided into K groups for K-fold validation in total. In each evaluation process, K-1 groups were used as the training set, and the remaining data was used as the validation set. The metrics of each process were recorded respectively. Finally, all the average metrics were taken as the result of cross validation. Independent test is mainly used to test the generalization ability of our model.

## 3 Results and discussion

### 3.1 Comparison with different feature representation combinations on different machine learning classifiers

#### Comparison on different machine learning classifiers

First, some experiments were conducted on cross-validation datasets and independent datasets to test performance of different machine learning classifiers, such as DT, KNN, SVM, LR, RF and lightGBM.

On the ACP740 dataset, Table 2 and Fig. 2 showed the performances of some different classifiers. The five metrics of the lightGBM classifier with the combined two-level features demonstrated the superior performance (Acc=0.897, Sp=0.918, Prec=0.917, F1=0.895, MCC=0.796, AUC=0.936). Only Sn=0.875 was lower than that from KNN. However, on the ACP240 dataset, the optimal value distribution of multiple metrics on different classifiers was relatively scattered, according to Table 3 and Fig. 3. The best results were achieved with the combined two-level features on DT with Acc=0.908, Prec=0.915 and F1=0.911, on RF with Sp=0.886 and AUC=0.947, on LightGBM with Sn=0.926 and MCC=0.800, respectively. Although lightGBM did not show consistent best performance, it could be seen from Table 3 that lightGBM achieved excellent results on Acc=0.900, Sp=0.875, Prec=0.895, and AUC=0.932. In addition, some different classifiers on the Main dataset and Alternate dataset were performed. On the Main dataset, it could be observed form Table 4 that lightGBM using combined two-level features also achieved consistent advantage as the best results (Acc=0.786, F1=0.802, MCC=0.579, AUC=0.834). However, on the Alternate dataset, Table 5 indicated that KNN achieved excellent performance. Most metrics were the maximum values (Acc=0.935, Sp=0.969, Prec=0.967, F1=0.933, MCC=0.872, AUC=0.971), but Sn=0.902 was less than the maximum value by 0.5%.

**Table 2.** Five-fold cross-validation performance comparison with different machine learning models on the ACP740 dataset.

| Features | Models | Acc | Sn | Sp | Prec | F1 | MCC | AUC |
| --- | --- | --- | --- | --- | --- | --- | --- | --- |
| peptide level feature | DT | 0.847 | 0.854 | 0.836 | 0.846 | 0.849 | 0.695 | 0.904 |
|  | KNN | 0.850 | 0.880 | 0.816 | 0.835 | 0.857 | 0.699 | 0.913 |
|  | SVM | 0.857 | 0.888 | 0.818 | 0.837 | 0.861 | 0.711 | 0.914 |
|  | LR | 0.841 | 0.848 | 0.823 | 0.844 | 0.844 | 0.680 | 0.895 |
|  | RF | 0.870 | 0.833 | 0.905 | 0.903 | 0.866 | 0.741 | 0.919 |
|  | lightGBM | 0.882 | 0.860 | 0.899 | 0.901 | 0.879 | 0.764 | 0.923 |
| residue level feature | DT | 0.838 | 0.872 | 0.797 | 0.822 | 0.844 | 0.679 | 0.896 |
|  | KNN | 0.827 | 0.843 | 0.806 | 0.819 | 0.830 | 0.653 | 0.891 |
|  | SVM | 0.831 | 0.844 | 0.820 | 0.830 | 0.835 | 0.666 | 0.880 |
|  | LR | 0.826 | 0.838 | 0.815 | 0.823 | 0.829 | 0.653 | 0.877 |
|  | RF | 0.847 | 0.873 | 0.822 | 0.835 | 0.852 | 0.697 | 0.902 |
|  | lightGBM | 0.851 | 0.879 | 0.819 | 0.836 | 0.856 | 0.702 | 0.906 |
| peptide level feature + residue level feature | DT | 0.858 | 0.862 | 0.855 | 0.861 | 0.859 | 0.720 | 0.920 |
|  | KNN | 0.878 | **0.891** | 0.865 | 0.868 | 0.879 | 0.756 | 0.924 |
|  | SVM | 0.869 | 0.883 | 0.856 | 0.862 | 0.872 | 0.740 | 0.916 |
|  | LR | 0.838 | 0.850 | 0.822 | 0.843 | 0.845 | 0.675 | 0.880 |
|  | RF | 0.889 | 0.874 | 0.907 | 0.906 | 0.889 | 0.779 | 0.929 |
|  | lightGBM | **0.897** | 0.875 | **0.918** | **0.917** | **0.895** | **0.796** | **0.936** |

**Fig.2.**
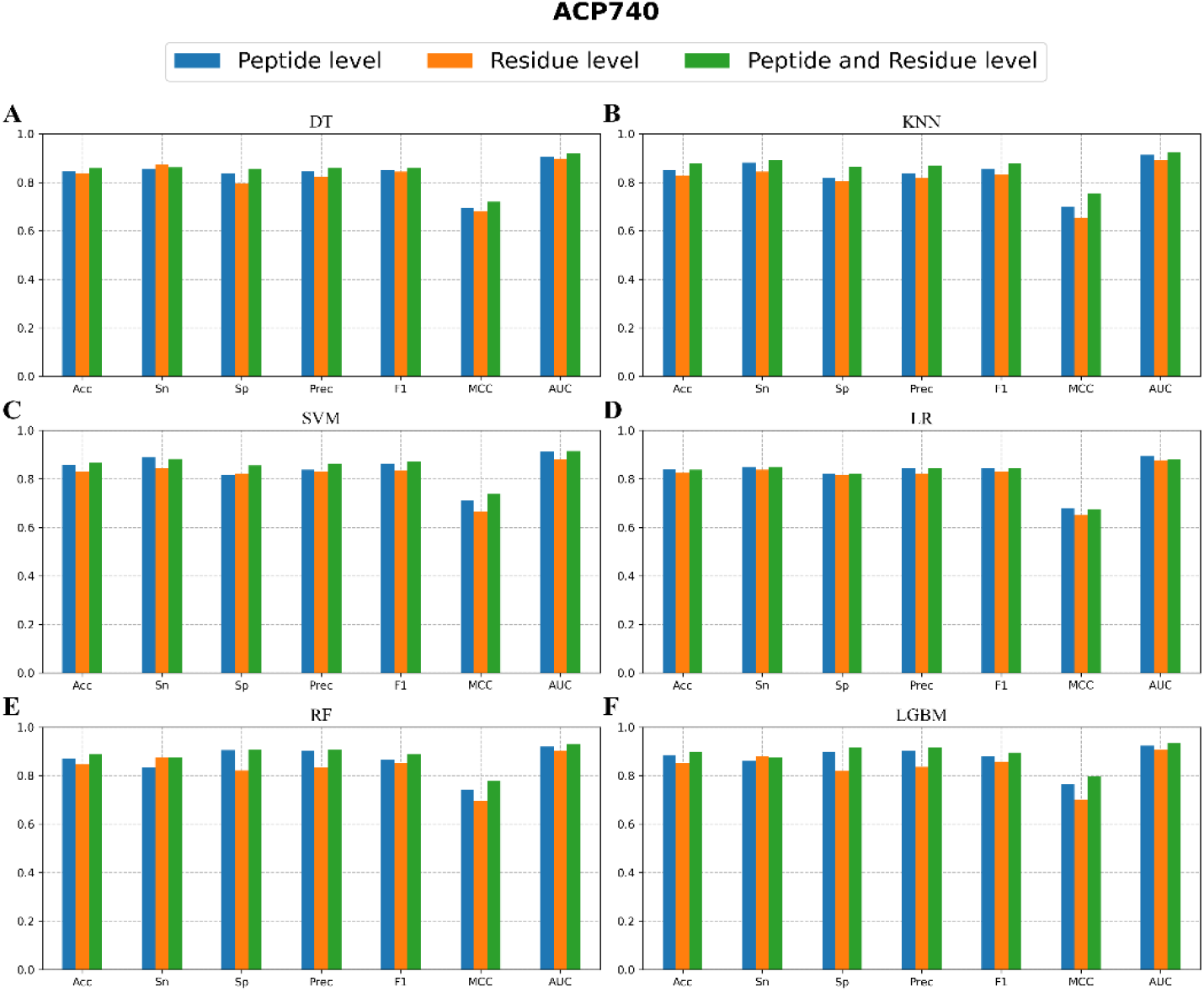
Five-fold cross-validation performance comparison of six classifiers with different feature combination on ACP740. The blue bars donated peptide level features, orange bars donated residue level features and green bars donated the combined two-level feature.

**Table 3.** Five-fold cross-validation performance comparison with different machine learning models on the ACP240 dataset.

| Features | Models | Acc | Sn | Sp | Prec | F1 | MCC | AUC |
| --- | --- | --- | --- | --- | --- | --- | --- | --- |
| peptide level<br>feature | DT | 0.783 | 0.739 | 0.839 | 0.849 | 0.783 | 0.584 | 0.822 |
|  | KNN | 0.746 | 0.885 | 0.544 | 0.693 | 0.775 | 0.462 | 0.787 |
|  | SVM | 0.758 | 0.867 | 0.530 | 0.726 | 0.784 | 0.451 | 0.816 |
|  | LR | 0.792 | 0.839 | 0.722 | 0.790 | 0.809 | 0.585 | 0.823 |
|  | RF | 0.775 | 0.778 | 0.796 | 0.834 | 0.784 | 0.586 | 0.831 |
|  | lightGBM | 0.783 | 0.814 | 0.725 | 0.805 | 0.799 | 0.565 | 0.831 |
| residue level<br>feature | DT | 0.904 | **0.956** | 0.851 | 0.886 | 0.915 | 0.818 | 0.931 |
|  | KNN | 0.787 | 0.902 | 0.666 | 0.764 | 0.821 | 0.595 | 0.861 |
|  | SVM | 0.854 | 0.861 | 0.865 | 0.877 | 0.863 | 0.721 | 0.850 |
|  | LR | 0.763 | 0.898 | 0.619 | 0.749 | 0.804 | 0.550 | 0.901 |
|  | RF | **0.908** | 0.931 | 0.881 | 0.904 | **0.916** | 0.815 | 0.946 |
|  | lightGBM | **0.908** | 0.925 | 0.870 | 0.909 | 0.915 | 0.796 | 0.929 |
| peptide level<br>feature +<br>residue level<br>feature | DT | **0.908** | 0.911 | 0.857 | **0.915** | 0.911 | 0.794 | 0.919 |
|  | KNN | 0.771 | 0.906 | 0.615 | 0.763 | 0.817 | 0.559 | 0.872 |
|  | SVM | 0.854 | 0.910 | 0.804 | 0.843 | 0.870 | 0.720 | 0.889 |
|  | LR | 0.817 | 0.916 | 0.708 | 0.781 | 0.841 | 0.640 | 0.876 |
|  | RF | 0.896 | 0.909 | **0.886** | 0.905 | 0.904 | 0.796 | **0.947** |
|  | lightGBM | 0.900 | 0.926 | 0.875 | 0.895 | 0.908 | **0.800** | 0.932 |

**Fig.3.**
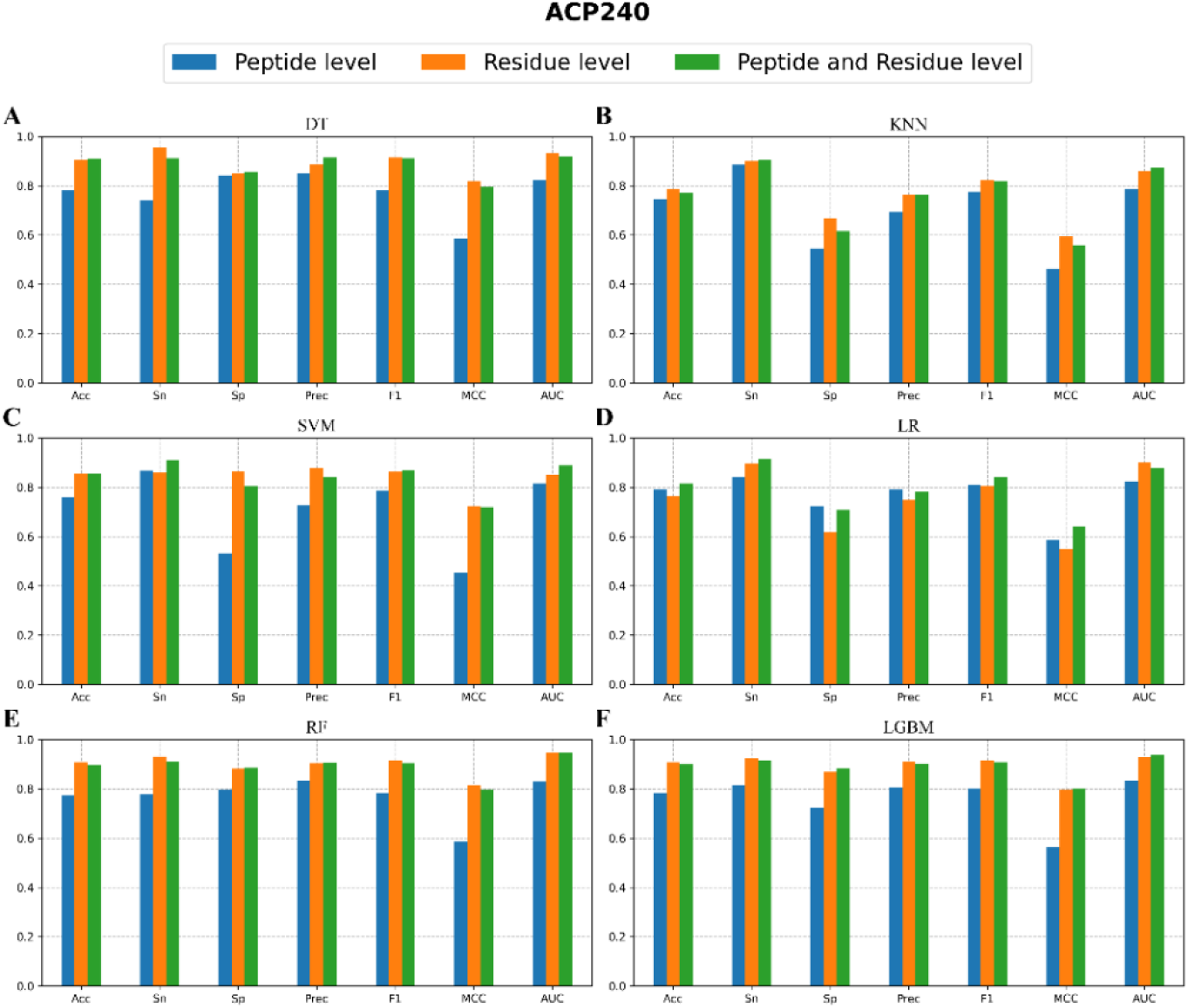
Five-fold cross-validation performance comparison of six classifiers with different feature combination on the ACP240. The blue bars donated peptide level features, orange bars donated residue level features and green bars donated both level feature.

**Table 4.** Independent test performance comparison with different machine learning models on Main dataset

| Features | Models | Acc | Sn | Sp | Prec | F1 | MCC | AUC |
| --- | --- | --- | --- | --- | --- | --- | --- | --- |
| peptide level<br>feature | DT | 0.768 | 0.678 | 0.859 | 0.829 | 0.746 | 0.546 | 0.811 |
|  | KNN | 0.727 | 0.532 | **0.924** | 0.875 | 0.662 | 0.495 | 0.819 |
|  | SVM | 0.754 | 0.585 | **0.924** | **0.885** | 0.704 | 0.540 | 0.824 |
|  | LR | 0.724 | 0.754 | 0.694 | 0.713 | 0.733 | 0.449 | 0.768 |
|  | RF | 0.771 | 0.725 | 0.818 | 0.800 | 0.761 | 0.545 | 0.811 |
|  | lightGBM | 0.783 | 0.749 | 0.818 | 0.805 | 0.776 | 0.567 | 0.826 |
| residue level<br>feature | DT | 0.733 | 0.801 | 0.665 | 0.706 | 0.751 | 0.470 | 0.764 |
|  | KNN | 0.707 | 0.731 | 0.682 | 0.698 | 0.714 | 0.414 | 0.739 |
|  | SVM | 0.730 | 0.801 | 0.659 | 0.703 | 0.749 | 0.465 | 0.786 |
|  | LR | 0.710 | 0.807 | 0.612 | 0.676 | 0.736 | 0.427 | 0.766 |
|  | RF | 0.754 | 0.830 | 0.676 | 0.721 | 0.772 | 0.513 | 0.798 |
|  | lightGBM | 0.760 | 0.813 | 0.706 | 0.735 | 0.772 | 0.522 | 0.802 |
| peptide level<br>feature +<br>residue level<br>feature | DT | 0.777 | 0.842 | 0.712 | 0.746 | 0.791 | 0.559 | 0.831 |
|  | KNN | 0.739 | **0.877** | 0.600 | 0.688 | 0.771 | 0.497 | 0.812 |
|  | SVM | 0.748 | 0.842 | 0.653 | 0.709 | 0.770 | 0.504 | 0.807 |
|  | LR | 0.748 | 0.772 | 0.724 | 0.737 | 0.754 | 0.496 | 0.828 |
|  | RF | 0.777 | 0.784 | 0.771 | 0.775 | 0.779 | 0.554 | 0.824 |
|  | lightGBM | **0.786** | 0.865 | 0.706 | 0.747 | **0.802** | **0.579** | **0.834** |

**Table 5.** Independent test performance comparison with different machine learning models on the Alternate dataset

| Features | Models | Acc | Sn | Sp | Prec | F1 | MCC | AUC |
| --- | --- | --- | --- | --- | --- | --- | --- | --- |
| peptide level<br>feature | DT | 0.907 | 0.886 | 0.927 | 0.924 | 0.905 | 0.814 | 0.961 |
|  | KNN | **0.935** | **0.917** | 0.953 | 0.952 | **0.934** | **0.871** | **0.969** |
|  | SVM | 0.922 | 0.855 | **0.990** | **0.988** | 0.917 | 0.852 | 0.961 |
|  | LR | 0.912 | 0.870 | 0.953 | 0.949 | 0.908 | 0.827 | 0.956 |
|  | RF | 0.925 | 0.860 | **0.990** | **0.988** | 0.920 | 0.857 | 0.967 |
|  | lightGBM | 0.933 | 0.902 | 0.964 | 0.961 | 0.930 | 0.867 | 0.966 |
| residue level<br>feature | DT | 0.847 | 0.803 | 0.891 | 0.881 | 0.840 | 0.697 | 0.908 |
|  | KNN | 0.855 | 0.850 | 0.860 | 0.859 | 0.854 | 0.710 | 0.910 |
|  | SVM | 0.837 | 0.829 | 0.845 | 0.842 | 0.836 | 0.674 | 0.895 |
|  | LR | 0.826 | 0.736 | 0.917 | 0.899 | 0.809 | 0.664 | 0.889 |
|  | RF | 0.870 | 0.808 | 0.933 | 0.923 | 0.862 | 0.747 | 0.920 |
|  | lightGBM | 0.870 | 0.855 | 0.886 | 0.882 | 0.868 | 0.741 | 0.917 |
| peptide level<br>feature +<br>residue level<br>feature | DT | 0.925 | 0.907 | 0.943 | 0.941 | 0.923 | 0.850 | 0.963 |
|  | KNN | **0.935** | 0.902 | 0.969 | 0.967 | 0.933 | **0.872** | **0.971** |
|  | SVM | 0.920 | 0.891 | 0.948 | 0.945 | 0.917 | 0.841 | 0.945 |
|  | LR | 0.907 | 0.881 | 0.933 | 0.929 | 0.904 | 0.815 | 0.928 |
|  | RF | 0.930 | 0.896 | 0.964 | 0.961 | 0.928 | 0.862 | 0.971 |
|  | lightGBM | 0.933 | 0.907 | 0.959 | 0.956 | 0.931 | 0.866 | 0.943 |

Although best results on four different datasets were achieved with different machine learning classifiers, the obvious phenomenon could be observed that lightGBM classifier obtained consistent satisfactory performance. On the ACP740 and the Main datasets, the lightGBM classifier outperformed other classifiers and obtained the preferable results on most metrics. The lightGBM classifier also was a satisfactory choice with the outstanding performance.

#### Comparison on different feature representation combinations

After appointing lightGBM as machine learning classifier in ME-ACP, multiple experiments also were conducted with peptide level and residue level features on cross-validation datasets and independent datasets to explore the effectiveness of combination with different level feature representations.

Fig. 2 and Table 2 showed the results of the five-fold cross-validation on the ACP740 dataset. The combined two-level features presented the superior performance compared with individual level feature. For example, the combined two-level features with the lightGBM classifier achieved the better performance (Acc=0.897, Sp=0.918, Prec=0.917, F1=0.895, MCC=0.796, AUC=0.936), except for Sn=0.875. On the ACP240 dataset, the combined two-level features with the lightGBM classifier also obtain the similar solution, judging from Fig. 3 and Table 3. Specifically, the four metrics (Sn=0.926, Sp=0.875, MCC=0.800, AUC=0.932) were better than the results from peptide level features or residue level features. Compared with maximum Acc=0.908, maximum Prec=0.909 and maximum F1=0.915, the combined two-level features still obtained a considerable performance (Acc=0.900, Prec=0.895, F1=0.908). Maybe overfitting occurred on small ACP240 dataset in HDNN. Similarly, multiple experiments were performed on the Main and the Alternate datasets. And the experimental results were shown in the Table 4 and Table 5, respectively. It could be seen that on the Main dataset, the results of the combined two-level features were generally better than the results of individual level feature in Table 4. For example, on the lightGBM classifier, five metrics have reached the optimal results (Acc=0.786, Sn=0.865, F1=0.802, MCC=0.579 and AUC=0.834). Only Sp=0.706 and Prec=0.747 is lower than the optimal value. Some optimal metrics values were observed at Table 5 (Acc=0.933, Sn=0.907, F1=0.931) on the Alternate dataset. But on the other three metrics (Sp=0.959, Prec=0.956, MCC=0.866, AUC=0.943), ME-ACP was just slightly lower than the optimal values.

In summary, by combining various feature extraction methods, our method obtained the superior performance. Moreover, the defects and shortcomings of different feature extraction methods are complemented mutually. Though using the probabilities from the machine learning models, our mothed also could convert high-dimensional protein features of different lengths into low-dimensional features of fixed length to reduce calculation cost in HDNN.

### 3.2 Comparison with different neural network structures

This section analyzed the influence on the performance with different DNN structures to identify ACPs and non-ACPs. According to the conclusion from the last section experiments, the combined two-level features and lightGBM were selected as the feature representations and machine learning classifiers respectively in this section to explore the effects of different DNNs.

For the DNN, three different deep neural networks were designed to find the optimal structure to improve the model performance on cross-validation datasets and the results were showed in Table 6 for ACP740 and ACP240. On the ACP740 dataset, except for Sn=0.875, the HDNN achieved the best performance (Acc=0.897, Sp=0.918, Prec=0.917, F1=0.895, MCC=0.796, AUC=0.936). On the other smaller dataset ACP240, the HDNN achieved the optimal effect only on Sp=0.875, Prec=0.895, AUC=0.932. Maybe overfitting occurred because of the small ACP240 dataset. From the experimental results of other two datasets, ME-ACP could achieve excellent effects on the larger datasets. The results of different DNNs on Independent datasets were showed in Table 7 for the Main dataset and the Alternate dataset. HDNN resulted in the improvement of most metrics (Acc=0.786, Sn=0.865, F1=0.802, MCC=0.579). Only Sp=0.706, Prec=0.747 and AUC=0.834 were lower than the maximum value. The Alternate dataset also achieved similar performance, Acc=0.933, Sn= 0.907, F1=0.931, MCC=0.866, achieving the superior results. Only Sp=0.959, Prec=0.956, AUC=0.943 were slightly lower than the optimal values.

**Table 6.** Five-fold cross-validation performance comparison with different DNNs which used the combined two-level feature representations on the cross-validation dataset.

| Datasets | DNN | Acc | Sn | Sp | Prec | F1 | MCC | AUC |
| --- | --- | --- | --- | --- | --- | --- | --- | --- |
| ACP740 | Residual | 0.864 | **0.901** | 0.822 | 0.842 | 0.870 | 0.728 | 0.916 |
|  | Bi-LSTM | 0.893 | 0.887 | 0.899 | 0.903 | 0.893 | 0.790 | 0.921 |
|  | residual + Bi-LSTM | **0.897** | 0.875 | **0.918** | **0.917** | **0.895** | **0.796** | **0.936** |
| ACP240 | residual | **0.908** | **0.956** | 0.858 | 0.888 | **0.918** | **0.821** | 0.930 |
|  | Bi-LSTM | 0.896 | 0.922 | 0.865 | 0.887 | 0.904 | 0.791 | 0.928 |
|  | residual + Bi-LSTM | 0.900 | 0.926 | **0.875** | **0.895** | 0.908 | 0.800 | **0.932** |

**Table 7.** Independent test performance comparison with different DNNs which used the combined two-level feature representations on independent dataset

| Datasets | DNN | Acc | Sn | Sp | Prec | F1 | MCC | AUC |
| --- | --- | --- | --- | --- | --- | --- | --- | --- |
| Main | residual | **0.786** | 0.854 | 0.718 | 0.753 | 0.800 | 0.577 | **0.837** |
|  | Bi-LSTM | 0.783 | 0.737 | **0.829** | **0.813** | 0.773 | 0.569 | 0.832 |
|  | residual + Bi-LSTM | **0.786** | **0.865** | 0.706 | 0.747 | **0.802** | **0.579** | 0.834 |
| Alt | residual | 0.925 | **0.907** | 0.943 | 0.941 | 0.923 | 0.850 | **0.964** |
|  | Bi-LSTM | 0.930 | 0.886 | **0.974** | **0.972** | 0.927 | 0.863 | 0.956 |
|  | residual + Bi-LSTM | **0.933** | **0.907** | 0.959 | 0.956 | **0.931** | **0.866** | 0.943 |

From a comprehensive perspective of various metrics, the results and the discusses demonstrated the advantage of HDNN compared with other DNNs, which support the efficiency of HDNN.

### 3.3 Comparison with different extraction length on residue level sequences

This section discussed the influence of the different sequence extraction length on residue level feature representation method on the ACPs and no-ACPs classification performance. The extraction length parameter *k* was set from initial length to maximum length. In order to make all protein sequences have the same feature dimension in each dataset, maximum length was set as the length of shortest common subsequence in each dataset. The maximum length in ACP740 ACP240, Main, Alternate data is 11, 11, 3, 3 respectively. For each residue level method, feature extraction started from the N-terminal or C-terminal of the protein sequence, respectively. To reduce feature redundancy, the optimal initial length varied from 1 to maximum length was selected based on performance on cross validation.

As shown in the Table 8, on the ACP740 dataset, the optimal values of metrics were not focused on ones. For example, Acc achieved the best result when the initial extraction length is 1 or 6 with values of 0.897, and Sn has the optimal value when length=10. However, on the ACP240 dataset, the overall better results appeared when length=6, and the best results were achieved on Acc=0.913, Sn=0.963, F1=0.921 and Acc=0.930 from Table 9. On the Main dataset, the best initial length is observed in Table 10 when length=3. It achieved the best results on Acc=0.792, Sp=0.835, Prec=0.821, MCC=0.586, AUC=0.855. In contrast, it was observed on the Alternate dataset in Table 11 that ME-ACP achieved excellent result when initial length was 3, with Acc=0.933, Sn=0.917, F1=0.932, AUC=0.970.

**Table 8.** Five-fold cross-validation metrices comparison of different sequence initial extraction length on residue level feature representation methods on ACP740 dataset.

| Initial length | Acc | Sn | Sp | Prec | F1 | MCC | AUC |
| --- | --- | --- | --- | --- | --- | --- | --- |
| 1 | **0.897** | 0.875 | 0.918 | 0.917 | 0.895 | **0.796** | 0.936 |
| 2 | 0.895 | 0.889 | 0.896 | 0.903 | 0.895 | 0.790 | **0.942** |
| 3 | 0.896 | 0.867 | 0.922 | 0.925 | 0.893 | **0.796** | 0.934 |
| 4 | 0.891 | 0.881 | 0.902 | 0.902 | 0.891 | 0.781 | 0.933 |
| 5 | 0.895 | 0.857 | **0.932** | **0.931** | 0.890 | 0.793 | 0.934 |
| 6 | **0.897** | 0.881 | 0.915 | 0.915 | **0.897** | 0.796 | 0.933 |
| 7 | 0.888 | 0.862 | 0.914 | 0.913 | 0.887 | 0.777 | 0.933 |
| 8 | 0.886 | 0.890 | 0.885 | 0.888 | 0.888 | 0.774 | 0.930 |
| 9 | 0.896 | 0.882 | 0.908 | 0.910 | 0.895 | 0.792 | 0.932 |
| 10 | 0.889 | **0.895** | 0.883 | 0.890 | 0.892 | 0.778 | 0.933 |
| 11 | 0.891 | 0.892 | 0.889 | 0.894 | 0.892 | 0.781 | 0.941 |

**Table 9.** Five-fold cross-validation metrices comparison of different sequence initial extraction length on residue level feature representation methods on ACP240 dataset.

| Initial length | Acc | Sn | Sp | Prec | F1 | MCC | AUC |
| --- | --- | --- | --- | --- | --- | --- | --- |
| 1 | 0.900 | 0.926 | 0.875 | 0.895 | 0.908 | 0.800 | 0.932 |
| 2 | 0.908 | 0.956 | 0.858 | 0.886 | 0.918 | **0.821** | 0.934 |
| 3 | 0.904 | 0.948 | 0.859 | 0.883 | 0.913 | 0.810 | 0.952 |
| 4 | 0.900 | 0.919 | 0.878 | 0.899 | 0.907 | 0.803 | 0.934 |
| 5 | 0.904 | 0.923 | **0.886** | **0.903** | 0.911 | 0.810 | 0.930 |
| 6 | **0.913** | **0.963** | 0.859 | 0.886 | **0.921** | 0.830 | 0.929 |
| 7 | 0.908 | 0.946 | 0.864 | 0.895 | 0.918 | 0.820 | 0.924 |
| 8 | 0.900 | 0.934 | 0.857 | 0.882 | 0.906 | 0.800 | 0.929 |
| 9 | 0.904 | 0.947 | 0.858 | 0.882 | 0.913 | 0.809 | **0.949** |
| 10 | 0.900 | 0.946 | 0.848 | 0.881 | 0.911 | 0.804 | 0.930 |
| 11 | 0.892 | 0.930 | 0.843 | 0.881 | 0.903 | 0.783 | 0.930 |

**Table 10.** Independent-test metrices comparison of different sequence initial extraction length on residue level feature representation methods on Main dataset.

| Initial length | Acc | Sn | Sp | Prec | F1 | MCC | AUC |
| --- | --- | --- | --- | --- | --- | --- | --- |
| 1 | 0.786 | **0.865** | 0.706 | 0.747 | **0.802** | 0.579 | 0.834 |
| 2 | **0.792** | 0.766 | 0.818 | 0.809 | 0.787 | 0.584 | 0.836 |
| 3 | **0.792** | 0.749 | **0.835** | **0.821** | 0.783 | **0.586** | **0.855** |

**Table 11.** Independent-test metrices comparison of different sequence initial extraction length on residue level feature representation methods on Alternate dataset.

| Initial length | Acc | Sn | Sp | Prec | F1 | MCC | AUC |
| --- | --- | --- | --- | --- | --- | --- | --- |
| 1 | **0.933** | 0.907 | 0.959 | 0.956 | 0.931 | 0.866 | 0.943 |
| 2 | **0.933** | 0.896 | **0.969** | **0.966** | 0.930 | **0.868** | 0.968 |
| 3 | **0.933** | **0.917** | 0.948 | 0.947 | **0.932** | 0.866 | **0.970** |

According to the discuss, initial length=6 achieved the best results consistently on the ACP740 and ACP240 datasets. Initial length=3 outperformed other condition on the Main and the Alternate datasets. Considered of the constraint from the initial length of independent datasets, ME-ACP selected 3 as the initial length for all datasets.

### 3.4 Comparison with existing methods

For further verifying the effectiveness of our method, we compared ME-ACP with the existing methods including ACP-DL, DeepACPpred, ACP-MHCNN on the cross-validation datasets and iACP, PEPred-Suite, ACPpred-Fuse, ACPred-FL, ACPred, AntiCP, AntiCP_2.0, iACP-DRLF on independent datasets in this work. It is noted that the combined two-level features were inputted lighuGBM classifier, and the HDNN was employed to integrate the local and global information and final predicter.

The comparisons were showed in Table 12 on the ACP740 and ACP240 dataset. ME-ACP obtained obvious improvements (Acc=0.896, Sp=0.922, Prec=0.925, F1=0.893, MCC=0.796) on ACP740 dataset, which were 3.6%~7.6% higher than the existing optimal metrics values, and only Sn=0.867 ranked second, was inferior to the best result of 88.9%. On the ACP240 dataset, our method also outperformed other methods. ME-ACP achieved the best results in Acc=0.904, Sn=0.948, Prec=0.883, F1=0.913 and MCC=0.810, including 2.1%~9.4% metrics improvement only with the 4 % reduction on Sp, which was due to the balance of Sp and Sn.

**Table 12.** Comparison of performance values from our method with other methods on ACP740 and ACP240 cross-validation dataset

| Dataset | Methods | Acc | Sn | Sp | Prec | F1 | MCC |
| --- | --- | --- | --- | --- | --- | --- | --- |
| ACP740 | ACP-DL | 0.815 | 0.826 | 0.806 | 0.824 |  | 0.631 |
|  | DeepACPpred | 0.85 | 0.853 | 0.85 | 0.856 | 0.85 | 0.706 |
|  | ACP-MHCNN | 0.86 | **0.889** | 0.831 | 0.844 |  | 0.72 |
|  | GRCI-Net | 0.8230 | 0.8355 | 0.8213 | 0.8351 |  | 0.6470 |
|  | ME-ACP | **0.896** | 0.867 | **0.922** | **0.925** | **0.893** | **0.796** |
| ACP240 | ACP-DL | 0.854 | 0.846 | **0.899** | 0.803 |  | 0.714 |
|  | DeepACPpred | 0.858 | 0.880 | 0.840 | 0.862 | 0.869 | 0.716 |
|  | ACP-MHCNN | 0.830 | 0.901 | 0.756 | 0.811 |  | 0.670 |
|  | GRCI-Net | 0.8750 | 0.8888 | 0.8840 | 0.8664 |  | 0.7531 |
|  | ME-ACP | **0.904** | **0.948** | 0.859 | **0.883** | **0.913** | **0.810** |

ME-ACP also was compared with other methods on the Main and Alternate datasets in Table 13. On the Main dataset, except for Sn, which was lower than the optimal value, most metrics reached the maximum values, which were improvement of >1.7% compared with the best results of other methods. In addition, our method maintained a consistent advantage on the Alternate dataset, especially at Acc=0.933 and MCC=0.866. Other metrics also achieved competitive values. Overall, ME-ACP have an advantage over other existing methods, especially on Acc, which indicate the efficiency of ME-ACP

**Table 13.** Comparison of performance with other methods on Main and Alternate independent dataset

| Dataset | Methods | Acc | Sn | Sp | MCC |
| --- | --- | --- | --- | --- | --- |
| Main | iACP | 0.551 | 0.779 | 0.322 | 0.110 |
|  | PEPred-Suite | 0.535 | 0.331 | 0.738 | 0.080 |
|  | ACPpred-Fuse | 0.689 | 0.692 | 0.686 | 0.380 |
|  | ACPpred-FL | 0.448 | 0.671 | 0.225 | -0.12 |
|  | ACPpred | 0.535 | 0.856 | 0.214 | 0.090 |
|  | AntiCP | 0.506 | **1.000** | 0.012 | 0.070 |
|  | AntiCP_2.0 | 0.754 | 0.775 | 0.734 | 0.510 |
|  | iACP-DRLF | 0.775 | 0.807 | 0.743 | 0.550 |
|  | ME-ACP | **0.792** | 0.749 | **0.835** | **0.586** |
| Alternate | iACP | 0.776 | 0.784 | 0.964 | 0.550 |
|  | PEPred-Suite | 0.575 | 0.402 | 0.747 | 0.160 |
|  | ACPpred-Fuse | 0.789 | 0.644 | 0.933 | 0.600 |
|  | ACPpred-FL | 0.438 | 0.602 | 0.256 | -0.15 |
|  | ACPpred | 0.853 | 0.871 | 0.835 | 0.710 |
|  | AntiCP | 0.900 | 0.897 | 0.902 | 0.800 |
|  | AntiCP_2.0 | 0.920 | **0.923** | 0.918 | 0.840 |
|  | iACP-DRLF | 0.930 | 0.896 | **0.964** | 0.860 |
|  | ME-ACP | **0.933** | 0.917 | 0.948 | **0.866** |

## 4 Conclusion

In this work, a novel method ME-ACP was proposed to identify ACPs and non-ACPs. In ME-ACP, residue level and peptide level features were fed into multiple lightGBM classifiers. And the concatenated probability vector from the classifiers were inputted into the HDNN for training. HDNN could integrate the local information and long-range dependency from the residue level and peptide level to discover further potential information.

The experimental results on cross-validation and independent datasets indicated ME-ACP achieved the competitive performances on the majority of metrics, especially on Acc. In addition, redundancy may exist in residue level feature from multiple feature extraction methods. Feature selection methods could be explored to find the optimal features in future work. Moreover, the architecture of ME-ACP could serve as a benchmark to adapt to the different tasks such as protein site prediction, antiviral peptide and protein homology detection etc.

## 5 Author contributions

GF and HY designed the overall scheme. RG, GF and CL developed the prediction models. RL, RH, and XF analyzed the data and results. GF and HY wrote the manuscript with input from all authors. RG and QM conceived the study and were in charge of overall direction and planning. All authors have read and approved the revised manuscript.

